# Abnormal Cannabidiol protects pancreatic beta cells in mouse models of experimental Type 1 diabetes

**DOI:** 10.1101/2020.12.16.423103

**Authors:** Isabel Gonzalez-Mariscal, Macarena Pozo Morales, Silvana Yanina Romero-Zerbo, Vanesa Espinosa-Jimenez, Alejandro Escamilla, Lourdes Sánchez-Salido, Nadia Cobo-Vuilleumier, Benoit R. Gauthier, Francisco Javier Bermudez-Silva

**Affiliations:** Instituto de Investigación Biomédica de Málaga-IBIMA, UGC Endocrinología y Nutrición. Hospital Regional Universitario de Málaga, Universidad de Málaga, 29009 Málaga, Spain; Microscopy platform, Biochemical Research Institute of Malaga (IBIMA), Malaga, Spain; Andalusian Center for Molecular Biology and Regenerative Medicine (CABIMER), Seville, Spain; Biomedical Research Center for Diabetes and Associated Metabolic Diseases (CIBERDEM), Madrid, Spain

**Keywords:** Abn-CBD, cannabinoids, type 1 diabetes, beta cell, insulitis, inflammation, T cells

## Abstract

**Background and Purpose:** The atypical cannabinoid Abn-CBD was reported to improve the inflammatory status in preclinical models of several pathologies including autoimmune diseases. However, its potential for autoimmune diabetes, *i*.*e*. type 1 diabetes (T1D), is unknown.

**Experimental Approach:** We used two mouse models of T1D, streptozotocin (STZ)-injected and non-obese diabetic (NOD) mice. Eight-to-ten-week-old male C57Bl6/J mice were pre-treated with Abn-CBD (1mg/kg of body weight) or vehicle for 1 week, following STZ treatment, and euthanized 1 week later. Six-week-old female NOD mice were treated with Abn-CBD (0.1-1mg/kg) or vehicle for 12 weeks and then euthanized. Blood, pancreas, pancreatic lymph nodes and circulating T cells were collected and processed for analysis. Glycemia was also monitored.

**Key Results:** Abn-CBD decreased circulating proinflammatory cytokines, ameliorated islet inflammation and the autoimmune attack, showing a 2-fold decrease in CD8^+^ T cells infiltration and reduced Th1/Th2 ratio in pancreatic lymph nodes of STZ-injected mice. Mechanistically, Abn-CBD reduced intra-islet phospho-NF-κB and TXNIP. Concomitant reduction of islet cell apoptosis and intra-islet fibrosis were observed in Abn-CBD pre-treated mice compared to vehicle. In NOD mice, Abn-CBD reduced the expression of *Ifng, Il21, Tnfa* and *Il10* while increased *Il4* in circulating CD4^+^ T cells compared to vehicle, reducing the severity of insulitis and improving glucose tolerance.

**Conclusion and Implications:** Altogether, we found that Abn-CBD reduces intra-islet inflammation and delays the progression of insulitis in mouse models of T1D, preserving healthy functional islets. Hence, Abn-CBD and related compounds emerge as new candidates to develop pharmacological strategies to treat early stages of T1D.

**WHAT IS ALREADY KNOWN:**

- Phytocannabinoids such as cannabidiol (CBD) have anti-inflammatory and glucose-lowering properties
- The CBD derivative Abn-CBD ameliorates inflammation in various diseases and modulates beta cell function

**WHAT THIS STUDY ADDS:**

- Abn-CBD reduces systemic and pancreatic inflammation in mice models of type 1 diabetes
- Abn-CBD prevents beta cell damage and loss during type 1 diabetes onset

**CLINICAL SIGNIFICANCE:**

- Synthetic cannabinoids emerge as potential treatment for type 1 diabetes

## INTRODUCTION

Glucose homeostasis is highly regulated by insulin, a hormone secreted by pancreatic beta cells necessary for the uptake, use and storage of glucose by targeted tissues. As such, insulin is a key hormone in glucose homeostasis and thus its secretion is a finely regulated process within the body. Sustained hyperglycemia, due to loss of insulin secretion and/or insulin action leads to diabetes. Insulin-dependent diabetes, known as type 1 diabetes (T1D), is an autoimmune disease that results in the progressive destruction of insulin-producing beta cells within pancreatic islets of Langerhans. Obliteration is preceded by the infiltration of immune cells in and around islets, a process known as insulitis (International Diabetes Federation).

Macrophages, CD4^+^ and CD8^+^ T cells play an important role in the development of T1D. T helpers (Th) cells, also known as CD4^+^ cells, upon activation and differentiation into specific T cell subsets, including helper type 1 (Th1) and 2 (Th2), Th17 and T regulatory (Treg) cells, are involved in the adaptive immune response by secreting specific sets of cytokines. An increase in the Th1/Th2 cell ratio is the tilting factor that initiates T1D (Talaat et al., 2016), while differentiation of Naïve CD4^+^ to Th17 cells is critical in the progression/pathogenesis of T1D (Alnek et al., 2015). Differentiation into Th1 cells is triggered by interferon gamma (IFNγ) and interleukin 12 (Il-12), whereas Th17 requires Il-6 and transforming growth factor beta (TGFβ). Both Th1 and Th17 T cells secrete pro-inflammatory cytokines (such as IFNγ, IL17 and tumor necrosis factor alpha -TNFα-) that induce intra-islet inflammation leading to the differentiation of CD8^+^ T cells to cytotoxic T lymphocytes (CTL). CTLs infiltrate the islets and induce beta cell death.

Despite the advances in our understanding of the factors involved in the pathogenesis, the prevalence of diabetes is on the rise, with an increase of 3% in juvenile diabetes in the last year (International Diabetes Federation). High blood glucose due to diabetes is associated with micro- and macro-vascular complications: atherosclerosis, strokes, heart attacks and heart failure and peripheral vascular disease (Forouhi and Wareham, 2014). Also, uncontrolled hyperglycemia increases the risk of renal failure, blindness, peripheral neuropathy and premature death (Forouhi and Wareham, 2014). Moreover, patients with T1D suffer from hypoglycemia, sometimes unnoticed, and of fatal consequences. To date no prevention or cure exists for T1D, hence patients require exogenous insulin injections for life. Current therapeutic approaches aim to reduce/prevent T-cell reactivity that is known to occur during the insulitis phase of T1D. However, the use of immunosuppressors such as anti-CD3 antibodies failed to show durable effects alone, and also they have non-islet, potentially life-threatening effects on the patient’s immune system (Couri et al., 2018). As such, alternative approaches are urgently needed to counteract T1D (Cobo-Vuilleumier and Gauthier, 2020).

Cannabinoids are known to regulate metabolism and immune action. While the roles of cannabinoids in T2D and obesity are fairly known (Engeli, 2008; Jourdan et al., 2013; Cinar et al., 2014; González-Mariscal et al., 2016a, 2016b, 2018), their effects on T1D have been largely unexplored. Cannabidiol (CBD) is a non-psychotropic phytocannabinoid from *Cannabis sativa* plant that has anti-inflammatory properties in a plethora of pathologies (Pacher et al., 2020), including diabetes (Weiss et al., 2008). Abnormal cannabidiol (Abn-CBD), a non-psychotropic synthetic cannabinoid, results from a transposition of the phenolic hydroxyl group and the pentyl side chain of CBD (Adams et al., 1977). Abn-CBD was initially described for its potent cardiovascular effects lowering blood pressure (Adams et al., 1977) and further attributed anti-tumoral (Tomko et al., 2019) and glucose management properties (McKillop et al., 2013; Ruz-Maldonado et al., 2018). In the pancreas, Abn-CBD has been reported to protect pancreatic beta cells from ER stress- and cytokine-induced apoptosis and enhance beta cell function in cultured islets of Langerhans and insulinoma cell lines (Ruz-Maldonado et al., 2018; Vong et al., 2019b, 2019a).

Here we aim to explore whether the atypical cannabinoid Abn-CBD can modulate the immune response during T1D onset in mice, and its potential as a therapeutic for the treatment of this disease in several mouse models of T1D.

## RESULTS

### Abn-CBD reduces streptozotocin-induced islet cells apoptosis without altering islet structure

Streptozotocin (STZ) is a widely used drug to mimic T1D, since it preferably gets internalized through GLUT2, induces reactive oxygen species (ROS) and provokes an inflammatory response that ultimately leads to beta cell death (Eleazu et al., 2013). As such we initially used this model to assess the impact of Abn-CBD on islet size and viability (Figure 1A). C57BL6J mice were pretreated with 1mg/kg of Abn-CBD or vehicle (saline:DMSO:Tween-80) for 7 days prior STZ treatment (Figure 1A). Mice were injected with BrdU 6 days after STZ injections and were then euthanized 24 h after (Figure 1A). As expected, STZ induced hyperglycemia. The prophylactic treatment with Abn-CBD slightly decreased glycemia without reaching statistical significance (390.5 ± 26.4 vs. 374.3 ± 29.4 mg/dl in vehicle- and Abn-CBD-treated mice, respectively; p=0.68). However, a significantly higher number of small islets correlating with a slightly lower average islet size was observed in the pancreas of Abn-CBD pre-treated mice as compared to vehicle-treated mice (Figure 1B-C). We did not find significant changes in intra-islet insulin staining (Figure 1D) nor in beta cell proliferation (Figure 1E-F). However, a significant 1.9-fold reduction in islet cell apoptosis was detected in Abn-CBD pretreated mice compared to vehicle, as shown by a reduction in intra-islet cleaved caspase 3 staining (Figure 1G-H). Thus, pretreatment with Abn-CBD prior STZ favors small islet size and blunts beta cell death.

**Figure 1.**
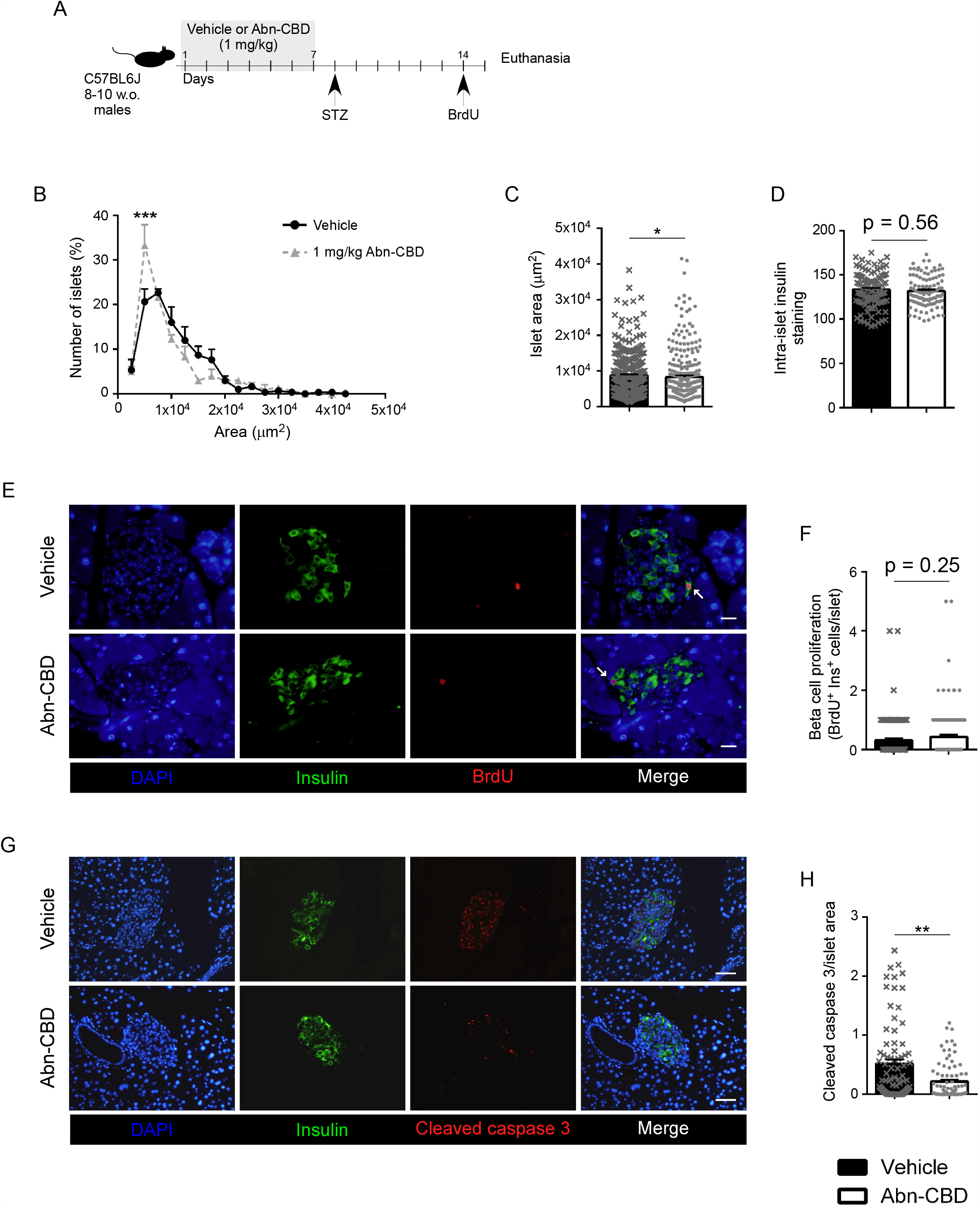
Effect of Abn-CBD on islet structure in STZ-induced T1D mice. (A) Schema of experimental procedure. Abn-CBD- and vehicle-treated mice were injected with streptozotocin (STZ), sacrificed one week later and pancreas dissected, formalin-fixed paraffin-embedded. Analysis of (B) islet area frequency distribution and (C) mean islet area per group. Pancreas sections were immunostained for insulin and (D) quantified. (E) Representative images of pancreas immunostained for BrdU (red), insulin (green) and DAPI (nuclei, blue) and (F) quantification of beta cell proliferation. (G) Representative images of pancreas immunostained for cleaved caspase 3 (red), insulin (green) and DAPI (blue) and (H) quantification of islet cell apoptosis. Data are mean ± S.E.M and single data points of vehicle- (black bars, grey crosses) and Abn-CBD-treated (white bars, grey dots) mice. Scale bar is 20 (E) and 50 µm (G). N = 9 mice per group, N = 90-100 islets per group. * p < 0.05, **p < 0.01 and ***p < 0.001 compared to vehicle. Non-significant p values are shown compared to vehicle.

### Abn-CBD attenuates STZ-induced apoptosis via the TXNIP and NF-ΚB pathways reducing systemic proinflammation

STZ was previously shown to induce the expression of thioredoxin-interacting protein (TXNIP), which is an activator of the inflammasome (Maedler et al., 2002; Zhou et al., 2010; Kelleher et al., 2014). In parallel, activation of the inflammasome by TXNIP also induces the phosphorylation of the p65 subunit of NF-κB resulting in its nuclear translocation and increased expression of proinflammatory cytokines which are then processed and activated by the inflammasome (Zhou et al., 2010; Kelleher et al., 2014). Together these pathways synergize in promoting beta cell death and fibrosis (Zhou et al., 2010; Artlett and Thacker, 2015). As such, we assessed whether Abn-CBD targeted TXNIP and NF-κB signaling pathways. Intra-islet TXNIP staining was significantly reduced (Figure 2A-B) with a concomitant 1.6-fold reduction in the activation of NF-κB in Abn-CBD-treated versus untreated mice (Figure 2C-D). Accordingly, fibrosis was reduced by 1.6-fold in islets from mice pre-treated with Abn-CBD as compared to vehicle-treated mice (Figure 2E-F). In parallel, circulating levels of the proinflammatory cytokines IL-6 and TNF alpha, as well as the chemokine CXCL-2 (MIP-2 alpha), were significantly reduced in Abn-CBD-treated mice as compared to vehicle-treated mice (Figure 2G-I). Interestingly, levels of IL-6 were undetectable in plasma of Abn-CBD treated mice. Although not statistically significant, MCP-1 and CXCL-1 were also decreased in mice treated with Abn-CBD (Figure 2J-K). Taken together these results indicate that Abn-CBD blunts systemic inflammation as well as enhances islet survival through amelioration of the TXNIP and NF-κB pathways.

**Figure 2.**
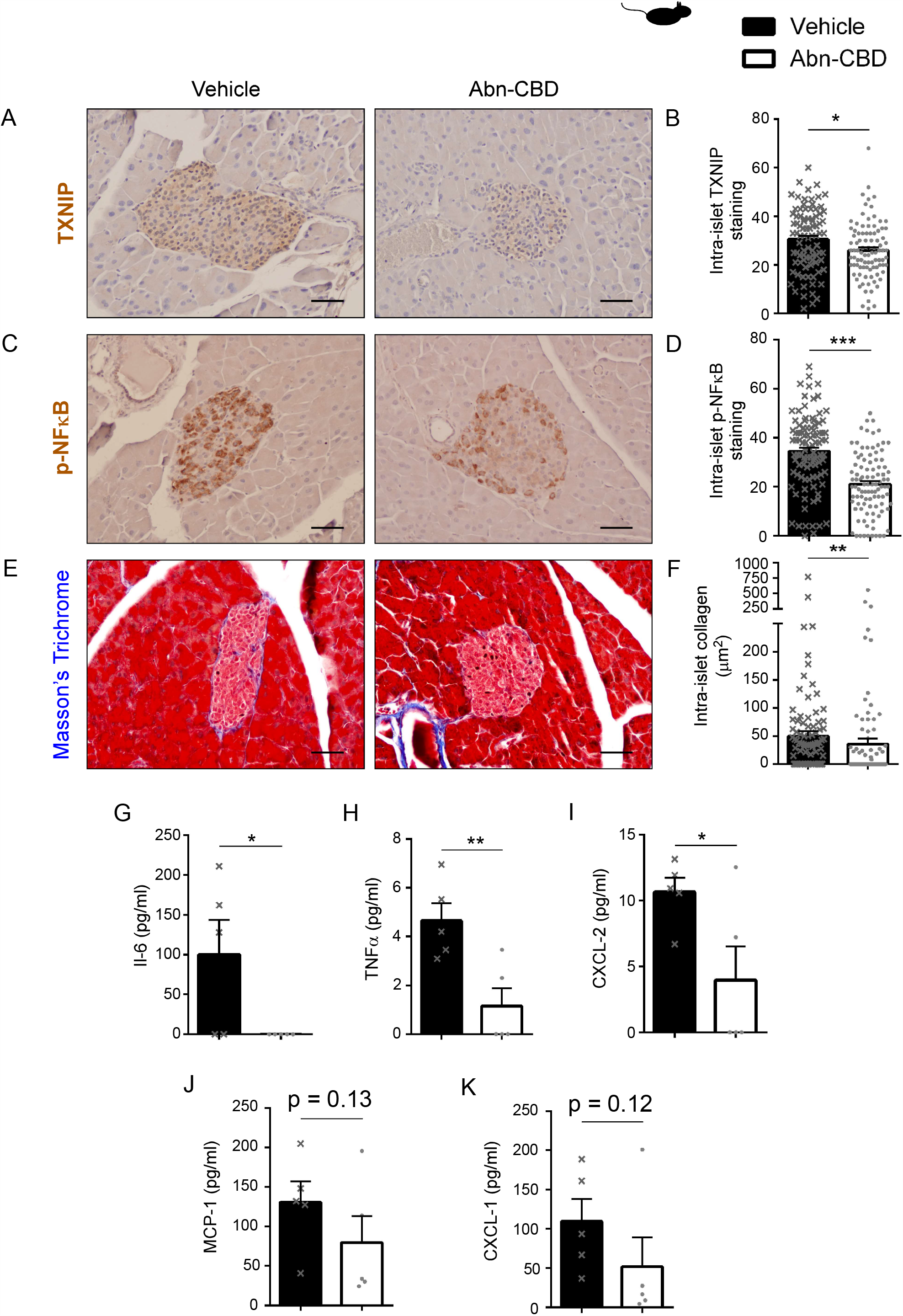
Effects of Abn-CBD on intra-islet and systemic inflammation in STZ-induced T1D mice. Representative images and quantification of pancreas sections immunstained for (A-B) TXNIP and (C-D) phospho-NFκB. (E) Representative images of pancreas sections stained to visualized the connective tissue (blue), nuclei (purple) and cytoplasm (pink) by Masson’s Trichrome staining. (F) Quantification by Image J FIJI of intra-islet collagen (blue staining) content upon image color deconvolution. Data are mean ± S.E.M and single data points of vehicle- (black bars, grey crosses) and Abn-CBD-treated (white bars, grey dots) mice. Scale bar is 20 µm. N = 9 mice per group, N = 90-100 islets per group. * p < 0.05, **p < 0.01 and ***p < 0.001 compared to vehicle. (G) Plasma levels of Interleukin 6 (IL-6), (H) tumor necrosis factor alpha (TNF-α), (I) chemokine ligand 2 (CXCL-2), (J) monocyte chemoattractant protein-1 (MCP-1) and (K) CXCL-1 in STZ-induced diabetic mice pretreated with Abn-CBD or vehicle. Data are mean ± S.E.M and single data points of vehicle- (black bars, grey crosses) and Abn-CBD-treated (white bars, grey dots) mice. N = 5 mice per group. * p < 0.05 and **p < 0.01 compared to vehicle. Non-significant p values are shown compared to vehicle.

### Abn-CBD decreases islet immune cell infiltration in STZ-treated mice

Since Abn-CBD dampens the expression of TXNIP and NF-κB activation, as well as circulating levels of proinflammatory chemo- and cytokines -indicative of inflammatory resolution-, we next assessed immune cell infiltration into the islets. Although no significant changes in the overall islet macrophage (F4/80^+^) population was detected (p=0.17; Figure 3A-B), the anti-inflammatory M2 subpopulations (CD163^+^) was slightly lower in Abn-CBD-treated compared to vehicle-treated mice (Figure 3C-D), and there was a significant 2.4-fold reduction in the number of proinflammatory M1 macrophages (iNOS^+^; Figure 3E-F). The number of T cells (CD3^+^) infiltrated into the islets was 1.8-fold lower in islets of mice pretreated with Abn-CBD (Figure 3G-H). Further analysis for T cell types showed a 2-fold reduction in the intra-islet number of CD8^+^ T cells in mice pre-treated with Abn-CBD compared to vehicle-treated mice (Figure 3I-J), while the overall T helper (Th, CD4^+^)-cell population remained constant (Figure 3K-L). However, treatment with Abn-CBD abolished the STZ-induced increase of the Th1/Th2 ratio (Figure 3M). Thus, Abn-CBD appears to blunt Th1-mediated activation and expansion of CTL cells correlating with lower islet cell apoptosis.

**Figure 3.**
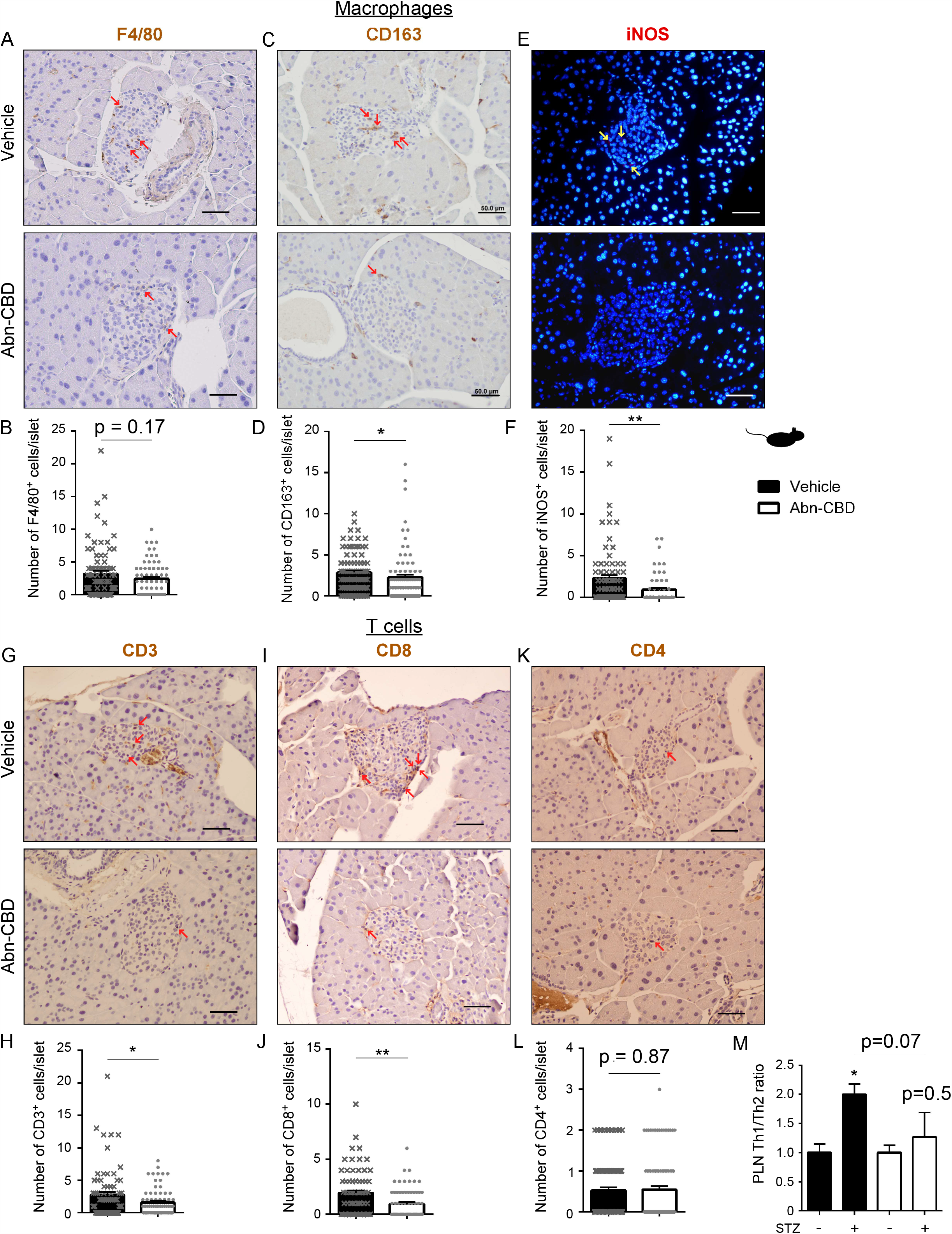
Analysis of immune cells infiltration in islets of Abn-CBD-treated STZ-induced T1D mice. Representative images and quantification of pancreas sections immunostained for (A-B) the macrophage marker F4/80, (C-D) the M2 polarized macrophage marker CD163 and (E-F) the M1 polarized macrophage marker iNOS, and (G-H) the T cell markers CD3, (I-J) CD4 and (K-L) CD8. Data are mean ± S.E.M and single data points of vehicle- (black bars, grey crosses) and Abn-CBD-treated (white bars, grey dots) mice. Scale bar is 50 µm. N = 9 mice per group, N = 90-100 islets per group. * p < 0.05 and **p < 0.01 compared to vehicle. Non-significant p values are shown compared to vehicle. (M) Ratio of Th1/Th2 cells in pancreatic lymph nodes (PLN). Data are mean ± S.E.M. N = 4 mice per group. * p < 0.05 compared to control (no STZ).

### Abn-CBD reduces the severity of insulitis in NOD mice

The non-obese diabetic (NOD) mouse is considered the gold standard model of T1D which is characterized by a spontaneous autoimmune attack predominantly in females and, eventually overt T1D within 20 weeks (Cobo-Vuilleumier et al., 2018). As such the impact of Abn-CBD was next assessed in this robust model. Six-week-old mice were treated i.p. with 0.1 or 1 mg/kg of Abn-CBD or vehicle for 12 weeks (Figure 4A). NOD mice treated with Abn-CBD showed a dose-dependent delay on the onset of T1D (onset of diabetes was considered when blood glucose > 250 mg/dl for 2 consecutive days) (Figure 4B). Upon 12 weeks, mice were sacrificed and pancreas analyzed by histochemistry for insulitis as per the following grades: grade 0 (no infiltration; healthy islets), grade 1 (immune cells approach the islet but do not infiltrate it; activation of the inflammatory response has started), grade 2 (immune cells infiltrate around the islet) and grade 3 (immune cells infiltrate into the islet; beta cell destruction occurs) (Figure 4C). The lower dose (0.1 mg/kg) of Abn-CBD reduced the progression of insulitis, increasing the number of islets in grade 2 insulitis (21 ± 4 vs. 11 ± 2%) while reducing grade 3 (42 ± 8 vs. 54 ± 10%) as compared to vehicle-treated mice (Figure 4D). A higher dose of Abn-CBD (1mg/kg) greater reduced the number of islets in grade 3 insulitis (36 ± 7 vs. 54 ± 10%), and importantly showed a significant increase of healthy (grade 0) islets (35 ± 8 vs. 25 ± 11%) (Figure 4D).

**Figure 4.**
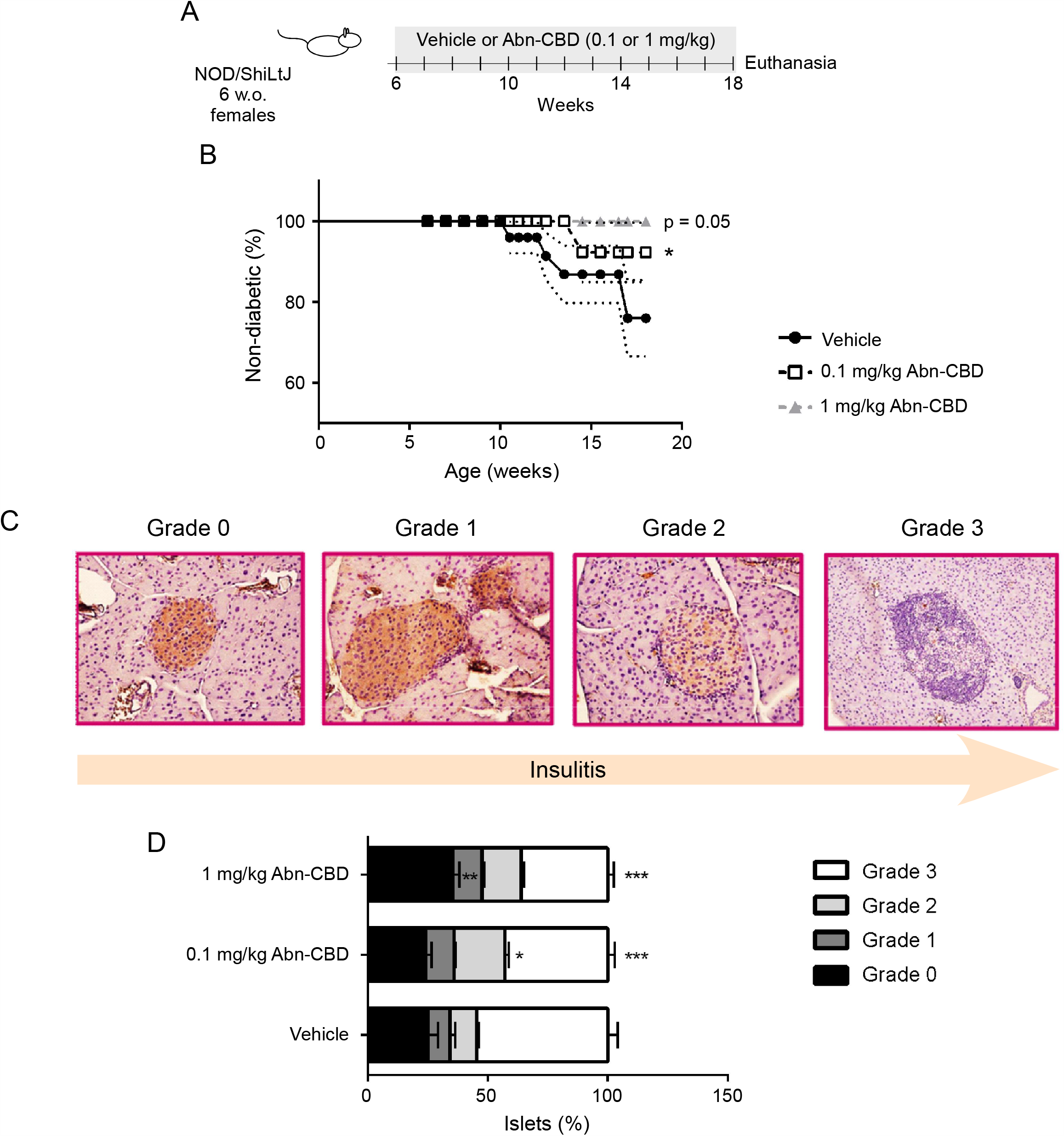
Diabetes incidence and insulitis grade in NOD mice treated with Abn-CBD. (A) Schema of experimental procedure. Six-week-old non-obese diabetic (NOD) mice were treated with Abn-CBD or vehicle for 3 months, and mice were sacrificed at 18-week-old and pancreas dissected, formalin-fixed and paraffin-embedded. Pancreas sections were immunostained for insulin and counterstained with hematoxylin. (B) Incidence of T1D in NOD mice treated with Abn-CBD or vehicle. Onset of diabetes was considered when blood glucose > 250 mg/dl for 2 consecutive days. (C) Distinct grades of insulitis were: grade 0 = no immune cell infiltration, insulin is preserved; grade 1 = immune cells approaching the islet, insulin is preserved; grade 2 = immune cells around the islet, islet start to lose insulin content; grade 3 = immune cells inside the islet, insulin is almost/totally lost. (D) Quantification of insulitis by determining the percentage of islets in each grade. N = 7-9 mice per group, N = 100 islets per group. N = 9-10 mice. * p < 0.05, **p < 0.01 and ***p < 0.001 compared to vehicle.

### Abn-CBD reduces activation of islet TXNIP-NF-κB pathway in NOD mice, ameliorating islet cells apoptosis

Treatment with 0.1 mg/kg of Abn-CBD reduced 1.5-fold islet cell apoptosis as compared to vehicle, as assessed by co-immunostaining of the pancreas with insulin and cleaved caspase 3 (Figure 5A-B). One mg/kg of Abn-CBD reduced islet cell apoptosis to the same extent as 0.1 mg/kg dose (Figure 5A-B). Accordingly, both dosages of Abn-CBD significantly reduced to the same extent TXNIP levels in islets of NOD mice (Figure 5C-D), and 1 mg/kg of Abn-CBD also induced a significant 1.5-fold reduction in the phosphorylation of NF-B (Figure 5E-F). Taken together, treatment of NOD mice with Abn-CBD significantly reduced islet inflammation, blunting the peri- and intra-islet infiltration of immune cells, thus improving islet survival.

**Figure 5.**
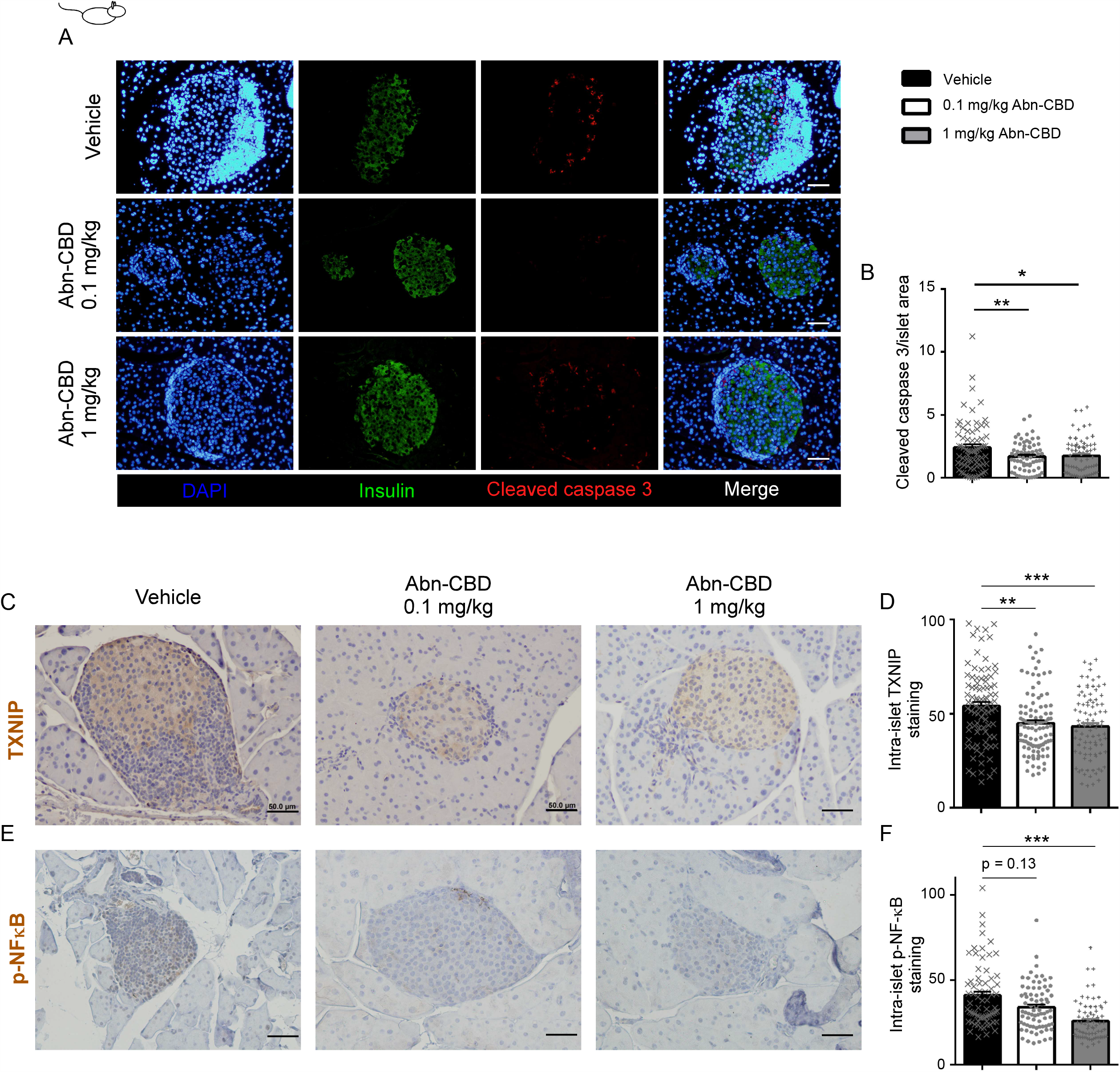
Apoptosis and inflammation of islet cells in NOD mice treated with Abn-CBD. (A) Representative images of pancreas of NOD mice treated with vehicle, 0.1 or 1 mg/kg of Abn-CBD immunostained for cleaved caspase 3 (red), insulin (green) and DAPI (blue) and (B) quantification of islet cell apoptosis. Representative images and quantification of pancreas sections immunstained for (C-D) TXNIP and (E-F) p-NF-κB. Data are mean ± S.E.M and single data points of vehicle- (black bars, grey crosses) and Abn-CBD-treated (white bars, grey dots) mice. Scale bar is 50 µm. N = 7-9 mice per group, N = 90-100 islets per group. * p < 0.05, **p < 0.01 and ***p < 0.001 compared to vehicle.

### Abn-CBD inhibits CD4^+^ Th1 cells in NOD mice

In the early stage of T1D, naїve CD4^+^ cells are polarized to Th1 T cells, increasing the Th1/Th2 ratio. Th1 CD4^+^ T cells secrete the pro-inflammatory cytokines IFNγ and TNFα that induce the differentiation of CD8^+^ T cells to CTL. CTLs infiltrate the islets of Langerhans and induce beta cell death by secreting granzymes and perforins. Since we found a significant reduction in CD8^+^ T cells infiltration into the islets of Abn-CBD-treated mice and decreased levels of systemic proinflammatory cytokines, we further analyzed the inflammatory profile of circulating Th cells. Circulating CD4^+^ T cells were isolated and the expression of cytokines mRNA analyzed. The purity of CD4^+^ T cells was determined by the expression of *Cd4* (Figure 6A). Treatment with Abn-CBD lowered the expression of *Il10* (T regulatory cell marker; Figure 6B), *Ifng* (Th1 cell marker; Figure 6C) and *Il21* (Th17 cells marker; Figure 6D), independently of the dose. CD4^+^ T cells from mice treated with 1 mg/kg Abn-CBD further showed significantly lower expression of *Tnfa* (Figure 6E). Expression of *Il4* was virtually absent in circulating CD4^+^ T cells from vehicle- or 0.1 mg/kg Abn-CBD-treated mice, while it was expressed in CD4^+^ T cells from 1 mg/kg Abn-CBD-treated mice (Figure 6F). Altogether, the data indicate a reduction in the Th1/Th2 ratio upon Abn-CBD treatment in NOD mice, similar to the decrease detected in Abn-CBD/STZ-treated mice (Figure 3M).

**Figure 6.**
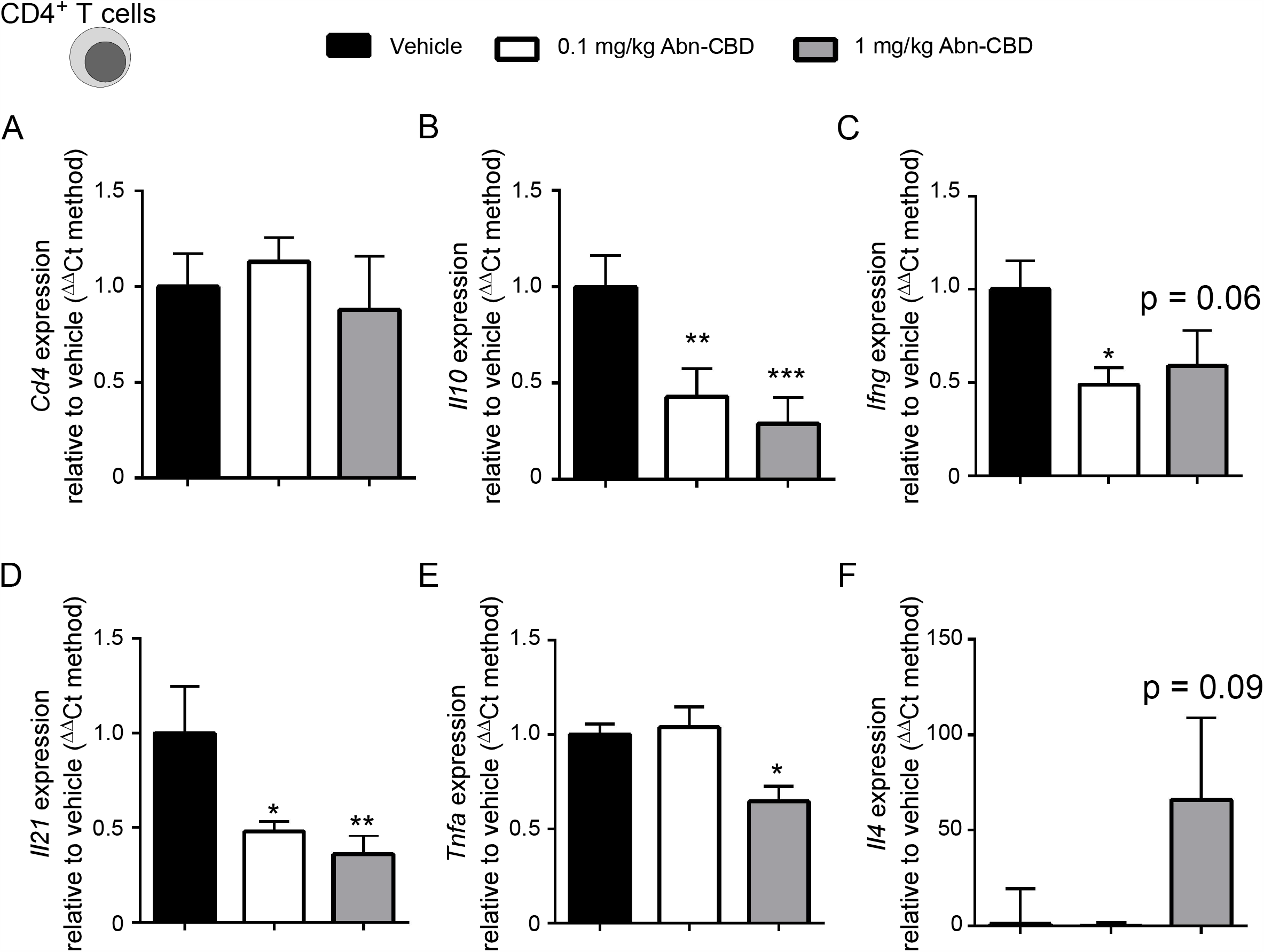
Expression profile of circulating CD4^+^ T cells in NOD mice treated with Abn-CBD. CD4^+^ T cells were isolated from NOD mice treated with vehicle, 0.1 or 1 mg/kg of Abn-CBD. (A) Expression of *Cd4* was used as control. Expression of (B) interleukin 10 (*Il10*), (C) interferon gamma (*Ifng*), (D) interleukin 21 (*Il21*), (E) *Tnfa* and (F) interleukin 4 (*Il4*) in CD4^+^ T cells. Data are mean ± S.E.M of vehicle-(black bars), 0.1 (grey bars) and 1 mg/kg (white bars) Abn-CBD-treated mice. N = 10 mice per group. * p < 0.05, **p < 0.01 and ***p < 0.001 compared to vehicle. Non-significant p values are shown compared to vehicle.

## DISCUSSION AND CONCUSIONS

Phytocannabinoids and synthetic cannabinoids have been investigated for their anti-inflammatory and glucose-lowering properties, but their potential as therapeutic agents for the treatment of insulitis and T1D remains unexplored. Herein we investigated the potential pre-clinical benefits of the synthetic cannabinoid Abn-CBD in T1D models, as emerging evidence point to its ability in modulating inflammation, insulin secretion as well as beta cell proliferation and apoptosis (McKillop et al., 2013, 2016; Ruz-Maldonado et al., 2018; Romero-Zerbo et al., 2020). Overall, we found that prophylactic administration of Abn-CBD was able to 1) reduce levels of systemic proinflammatory cytokines, 2) reduce polarization of CD4^+^ T cells towards proinflammatory Th1 cells, 3) decrease immune infiltration in the islets, the severity of insulitis and intra-islet inflammation and 4) decrease apoptosis of islet cells. These actions of Abn-CBD are compatible with an improved intra-islet environment potentially preserving islet function.

Prophylactic treatment with Abn-CBD slightly delayed the onset of T1D in NOD mice and decreased insulitis and fibrosis in NOD and STZ-injected mice, respectively, suggesting an improved intra-islet environment. Indeed, our finding of a higher percentage of small islets in Abn-CBD treated mice also suggests that Abn-CBD may be specifically protecting these small islets as they are known to be more sensitive to damage during type 1 diabetes (Tomita, 2010).

Abn-CBD significantly reduced beta cell apoptosis in both T1D mouse models, in agreement with previous reports in mouse models of type 2 diabetes (Ruz-Maldonado et al., 2018; Vong et al., 2019b; Romero-Zerbo et al., 2020), in which Abn-CBD was shown to reduce cytokine (mix of TNFα, IFNγ and IL-1β)-induced apoptosis in islets *in vitro* (Ruz-Maldonado et al., 2018). IL-6, which is elevated in T1D (Alnek et al., 2015), plays a very important role at the onset of T1D by activating CD4^+^ T cell differentiation (Hundhausen et al., 2016). Abn-CBD greatly reduced circulating levels of proinflammatory cytokines, including IL-6, whose levels were, in fact, undetectable. Accordingly, CD4^+^ T cells from Abn-CBD-treated NOD mice expressed significantly less *Tnfa* and *Ifng* than vehicle-treated NOD mice. Moreover, expression levels of *Il21*, a cytokine secreted by Th17 were also decreased in NOD mice. Th17 cells have been considered a central player in T1D development, which enhances the autoimmune response that leads to insulitis by further stimulating CD8^+^ cells differentiation into CTL cells (Alnek et al., 2015). Thus, decreased *Il21* expression further supports the establishment of an anti-inflammatory environment. This mechanism was also observed in the STZ model, in which Abn-CBD-treated mice also harbored a decreased number of T cell infiltration into the islets as well as a lower Th1/Th2 cell ratio, both involved in CTL cells expansion. We also observed decreased expression of *Il10*, which is secreted by Tregs, in addition to Th2 cells. Changes in *Il10* may be attributed to a modulation of Tregs, but little is known on the role of Tregs in T1D. Defects in Treg function are associated with T1D (Lindley et al., 2005), although no changes in Treg population have been found in patients with T1D (Brusko et al., 2007). Further studies regarding the potency of Abn-CBD to regulate these different T cell populations are necessary.

Macrophages can further amplify the inflammatory response by recruiting other immune cells such as lymphocytes. We did not detect changes in the total amount of macrophages in the islets of Abn-CBD-*versus* vehicle-treated STZ-injected mice; however, the number of intra-islets proinflammatory M1 macrophages, also known as classically activated macrophages, were significantly lower in the Abn-CBD-treated mice. IFN-γ induces the polarization of macrophages to M1 (Yao et al., 2019), which are involved in the Th1 response. Indeed, M1 macrophages secrete pro-inflammatory cytokines such as TNF-α and IL-6 (Yao et al., 2019), which were decreased in Abn-CBD-treated mice, as well as the Th1/Th2 ratio, compared to vehicle-treated mice.

We also investigated the molecular mechanisms underlying these anti-inflammatory and anti-apoptotic effects of Abn-CBD in the islets of T1D mice. Key components regulating inflammation and beta cell survival are the NF-κB pathway and the inflammasome which, when activated, synergistically work to promote beta cell death (Zhou et al., 2010; Artlett and Thacker, 2015). STZ-induced beta cell damage implicate elevated TXNIP expression and subsequent activation of the inflammasome (Maedler et al., 2002; Zhou et al., 2010; Kelleher et al., 2014). At the molecular level, we found that Abn-CBD repressed activation of both NF-κB (measured as phosphorylation of p65) and the inflammasome (measured as TXNIP staining) in islets from both models of T1D treated with Abn-CBD, that explains a reduced inflammation-mediated beta cell death and fibrosis. Thus, the anti-apoptotic effect of Abn-CBD in beta cells seems to arise from a combination of lower levels of cytokines and chemokines in circulation, decreased inflammation and immune infiltration into the islets, and reduced sensitivity to cytokines by beta cells (Ruz-Maldonado et al., 2018; Vong et al., 2019b; Romero-Zerbo et al., 2020) and this study. The reduced insulitis originates from lower CD4^+^ T cells polarization to Th1, dampening Th1/Th2 ratio, subsequently decreasing CTL cells maturation and infiltration and lowering the presence of M1 macrophages in the islets. On the other side, down-regulation of the NF-κB pathway and the inflammasome leads to reduced beta cell sensitivity to cytokines and reduced islet cytokines secretion, further preventing T cell infiltration.

We did not find a significant increase in beta cell proliferation in Abn-CBD-treated STZ-induced diabetic mice as compared to vehicle-treated mice that may account for the lack of improved glycemia over the short experimental period time. These results challenge previous reports indicating that Abn-CBD stimulated beta cell proliferation in the STZ-treated Swiss mouse model as well as in a diet-induced mouse model of obesity and prediabetes and isolated human islets (McKillop et al., 2016; Ruz-Maldonado et al., 2018; Romero-Zerbo et al., 2020). Discrepancies may potentially be related to differences in time and via of administration. McKillop et *al*. administered Abn-CBD orally and continued the treatment post-STZ administration for up to 30 days, while our Abn-CBD treatment was prophylactic, *i*.*e*. only administered before diabetes induction. Also, we delivered Abn-CBD intraperitoneally and not orally. Abn-CBD has been shown to have incretin-dependent effects (McKillop et al., 2016). Incretins, especially GLP-1, enhance beta cell proliferation (Doyle and Egan, 2007). Probably, the increased beta cell proliferation in orally-administered Abn-CBD-treated mice is an incretin-mediated effect. Similarly, the Abn-CBD-induced beta cell proliferation that we found in a diet-induced mouse model of obesity and prediabetes (Romero-Zerbo et al., 2020), could be an incretin-mediated effect as, though we injected Abn-CBD intraperitoneally, incretin levels have been reported to be up-regulated in obesity (Chia et al., 2017). However, we did not observe an impaired beta cell proliferation as it occurs with immunosuppressants (Nir et al., 2007). Consequently, Abn-CBD presents an advantage to immunosuppressants, since Abn-CBD would not avoid the recovery of the beta cell mass in T1D while it still lessened the insulitis.

In sum, our work in mouse models of T1D shows that Abn-CBD is a potent immunomodulator of the aberrant immune response characteristic of this disease. At diagnosis, around 50% of patients with T1D go through what is called the honeymoon phase for as long as 1 year, in which the residual beta cells can maintain normoglycemia without the need of exogenous insulin (Katchan et al., 2016). We herein provide evidence that cannabinoids such as the Abn-CBD modulate the immune response and beta cell death at the early stages of T1D. Hence, they are promising compounds that merit further clinical development and, if successful, they could be useful for pharmacological interventions aimed at stopping insulitis and beta cell loss at early stages of T1D, especially in patients entering the honeymoon phase of the disease.

## MATERIALS AND METHODS

### Materials

Abn-CBD was purchased from Tocris (Biogen). Streptozotocin (STZ), 5-Bromo-2’-deoxyuridine (BrdU) and Fluoroshield mounting media with DAPI were purchased form Sigma Aldrich. Antigen Unmasking Solution (H-3300) and DAB Peroxidase Substrate were purchased from Vector Laboratories. Trichrome Stain Kit was purchased from Abcam. Primary antibodies used were mouse anti-BrdU (1:50; G1G2) from the Hybridoma Bank, mouse anti-insulin (1:500; I2018) from Sigma Aldrich, rabbit anti-insulin (1:100; sc-9168) from Santa Cruz Biotechnology, rabbit anti-cleaved caspase 3 (1:100; #9661) from Cell Signaling, rat anti-F4/80 (1:100; ab6640) and rabbit anti-TXNIP (1:100; ab188865), anti-NF-kB p65 (phospho S536; 1:100; ab86299), anti-CD3 (1:100; ab5690), anti-CD163 (1:500; ab182422) and anti-iNOS (1:100; Ab15323) from Abcam, and mouse anti-CD4 and rabbit anti-CD8 from Ventana. Secondary antibodies were anti-mouse Alexa Fluor 488, anti-mouse Alexa Fluor 568, anti-rabbit Alexa Fluor 488 and rabbit Alexa Fluor 568 (1:1000; Thermo Fisher Scientific).

### Animals

Animal care and procedures were approved by the Animal Experimentation Ethics Committee of the Malaga University and authorized by the government of Andalucia (Project number 28/06/2018/107). Mice were housed in groups of 10 using 12 h dark/light cycles and provided regular chow (SAFE A04, Panlab) and water *ad libitum*. Eight-to-ten-week-old C57BL6J male and 6-week-old NOD/ShiLtJ female mice were purchased from Charles River France and Italy respectively. After 1 week of acclimation, C57BL6J male littermate mice were randomly assigned to 2 groups: vehicle or Abn-CBD. Mice were injected daily with intraperitoneal (i.p.) injections of vehicle (saline:DMSO:Tween-80 95:4:1) or 1 mg/kg of Abn-CBD for 7 days. Mice were then fasted for 4 h and then injected i.p. with 150 mg/kg of streptozotocin (STZ) and given 10% sucrose water for 48 h to avoid hypoglycemia. Blood glucose was monitored daily using OneTouch Ultra blood glucose meter (LifeScan IP Holdings, LLC). After 6 days mice were injected with BrdU and euthanized by cervical dislocation the following day. NOD/ShiLtJ female littermate mice were randomly assigned to 3 groups: vehicle, 0.1 or 1 mg/kg of Abn-CBD. Mice were daily i.p. injected with vehicle or different dose of Abn-CBD for 3 months (20-22-week-old). At the end of the study, mice were euthanized by cervical dislocation, and tissues and blood were collected and processed immediately for histological and biochemical analysis.

### Histopathology of pancreas

Pancreas were dissected and fixed in methanol-free 4% paraformaldehyde for 24 h at 4°C, then paraffin-embedded. For immunohistochemistry, 5 µm sections were dewaxed, rehydrated and subjected to heat-mediated citric acid-based antigen retrieval for 20 min in a pre-heated steamer and further 20 min of cooling down. For DAB staining, peroxidase activity was blocked for 10 min with 3% hydrogen peroxide. Sections were blocked in 5% goat or donkey serum or bovine serum albumin (BSA) 0.3% Triton-X-100 in PBS for 30 min at 37°C. Slides were then incubated with primary antibody diluted in 1% serum or BSA, 0.3% Triton-X-100 in PBS overnight at 4°C, washed 3 times for 5 minutes in 0.3% Triton-X-100 PBS, incubated with secondary antibody diluted in 1% serum or BSA, 0.3% Triton-X-100 in PBS for 30 min at 37°C and then washed 3 times for 5 minutes in 0.3% Triton-X-100 PBS. Alexa Fluor or HRP polymer-conjugated antibodies were incubated for 45 min at 37°C, and nuclei stained using DAPI for immunofluorescence. DAB staining was performed following hematoxylin for nuclei staining. Negative controls were performed using 0.3% Triton-X-100 PBS containing 1% goat serum or BSA. For Masson’s trichrome staining, 5 µm sections were dewaxed, rehydrated and stained using Trichrome Staining Kit following manufacturer’s instructions. Imaging was performed at 200X using an Olympus BX41. BrdU and insulin co-staining images were obtained using a Leica SP5 II confocal microscope. Densitometric analysis was performed using ImageJ (NIH). The percentage of proliferating beta-cells per islet was determined by counting the number of BrdU-insulin-positive cells. DAB staining was quantified using ImageJ Fiji after color deconvolution and further processed by Image J software to quantify signal intensity. Images from trichrome staining were color deconvoluted and area of islets with blue (collagen) signal quantified by Image J software. Infiltration of immune cells was quantified by counting the number of positive stained cells per islet.

### Flow cytometry

Pancreatic lymph nodes (PLN) were dissected and mechanically disaggregated in media (RPMI). Erythrocytes were lysated by incubating cells with ACK lysis buffer for 3 min at room temperature. Cells were immediately washed 3 times with media and plated at a density of 4×10^6^ cells/ml in a 96-well plate. Cells were stained using PerCP anti-mouse CD3 and APC anti-mouse CD4 (Biolegend) for 20 min at room temperature. Cells were washed with FACS Flow, fixed in 4% paraformaldehyde and washed with Perm/Wash Buffer (BD Biosciences) before staining with APCCy7 anti-mouse IFNγ (Biolegend). Appropriate isotype controls were used in all the experiments. Populations were determined by using BD FACS Canto II and the FACS Diva software (BD Biosciences).

### Hormones and cytokines measurements

Blood samples were collected in tubes containing EDTA and plasma separated by centrifugation. Plasma samples were flash frozen and kept at -80°C. Insulin was determined by ELISA (Mercodia), and cytokines were determined using a ProcartaPlex Immunoassay (Thermo Fisher Scientific).

### CD4^+^ T cell isolation

Blood samples from final bleed were collected in tubes containing heparin. CD4^+^ T cells were isolated using EasySep Mouse CD4+ T Cell Isolation Kit (Stemcell technologies) following manufacturer’s instructions. Briefly, red blood cells lysis was achieved with 0.8% ammonium chloride. Cells were washed with Ca^+2^-Mg^+2^-free PBS containing 2% fetal bovine serum (FBS) and 1 mM EDTA and resuspended at 1×10^8^ cells/ml. Samples were incubated with rat serum and isolation cocktail (1:40) for 10 min at room temperature. Then, samples were incubated with streptavidin-coated magnetic particles (RapidSpheres) for 2.5 min at room temperature and negative isolation performed using an EasySep magnet (#18000, Stemcell technologies). Isolated CD4^+^ T cells were pelleted and flash frozen.

### RNA purification and real time PCR

Total RNAs from isolated CD4^+^ T cells were obtained using TRI Reagent (Sigma Aldrich). Total RNA concentration and quality were measured by Nanodrop (Thermo Fisher Scientific). Reverse transcription was performed using SuperScript IV Reverse Transcriptase (Thermo Fisher Scientific). Relative expression of *Cd4, Tnfa, Ifng, Il4, Il10, Il17* and *Il21* genes was assayed using TaqMan Fast Advanced Master Mix and FAM-labeled TaqMan Gene Expression Assays for each specific gene (Thermo Fisher Scientific) on an ABI 7500 real time PCR System (Thermo Fisher Scientific). Duplex reactions were performed using VIC-labeled β-actin for endogenous control.

### Statistical analysis

Data are shown as mean ± SEM, including individual values. Statistical analysis was performed using GraphPad Prism version 6.07. Normal distribution of data was assessed by normality tests. Mean values were compared using Student’s t-test or Mann-Whitney test for two groups comparisons, and ANOVA with Tukey’s or Dunn’s test for multiple comparisons, for parametric or non-parametric test, respectively. Survival curves were compared using the two-group Mantel-Cox test. A p-value < 0.05 was considered significant.

### Randomization

Littermates were randomly assigned to vehicle or treatment groups.

### Blinding

Samples were coded and analysis of plasma and images were blinded to the analyst.

## Funding

H2020-MSCA-IF-2016, Grant Agreement number: 748749, EU. Consejeria de Salud y Familias, Junta de Andalucia (PI-0318-2018). Instituto de Salud Carlos III, Ministerio de Sanidad, Gobierno de España (PI17/01004). Ministerio de Ciencia, Innovación y Universidades, Agencia Estatal de Investigación and Fondo Europeo de Desarrollo Regional (BFU2017-83588-P TO BRG). We thank all the staff of the animal facility at IBIMA (ECAI de Experimentación Animal) and especially its coordinator Dr. Ricardo González Carrascosa for his excellent work. The authors also gratefully acknowledge all the staff of the bioimaging facility at IBIMA (ECAI de Imagen) and the proteomic (ECAI de Proteómica) facility, especially Dra. Carolina Lobo-García for excellent technical assistance.. FJBS and IGM belongs to the regional “Nicolás Monardes” research program from Consejería de Salud (C-0070-2012, RC0005-2016 and C1-0018-2019; Junta de Andalucía, Spain). FJBS, NCV and BRG are members of the pancreatic islets study group from the Spanish Society for Diabetes (SED). CIBERDEM is an initiative of the Instituto de Salud Carlos III.

## CONFLICT OF INTEREST

Authors declare no conflict of interest

## Notes

### Competing Interest Statement

The authors have declared no competing interest.

## REFERENCES

Adams, M.D., Earnhardt, J.T., Martin, B.R., Harris, L.S., Dewey, W.L., and Razdan, R.K. (1977). A cannabinoid with cardiovascular activity but no overt behavioral effects. Experientia 33: 1204– 5.

Alnek, K., Kisand, K., Heilman, K., Peet, A., Varik, K., and Uibo, R. (2015). Increased Blood Levels of Growth Factors, Proinflammatory Cytokines, and Th17 Cytokines in Patients with Newly Diagnosed Type 1 Diabetes. PLoS One 10: e0142976.

Artlett, C.M., and Thacker, J.D. (2015). Molecular Activation of the NLRP3 Inflammasome in Fibrosis: Common Threads Linking Divergent Fibrogenic Diseases. Antioxid. Redox Signal. 22: 1162–1175.

Brusko, T., Wasserfall, C., McGrail, K., Schatz, R., Viener, H.L., Schatz, D., et al. (2007). No Alterations in the Frequency of FOXP3+ Regulatory T-Cells in Type 1 Diabetes. Diabetes 56: 604–612.

Chia, C.W., Carlson, O.D., Liu, D.D., González-Mariscal, I., Santa-Cruz Calvo, S., and Egan, J.M. (2017). Incretin Secretion in Humans is under the Influence of Cannabinoid Receptors. Am. J. Physiol. -Endocrinol. Metab. ajpendo.00080.2017.

Cinar, R., Godlewski, G., Liu, J., Tam, J., Jourdan, T., Mukhopadhyay, B., et al. (2014). Hepatic cannabinoid-1 receptors mediate diet-induced insulin resistance by increasing de novo synthesis of long-chain ceramides. Hepatology 59: 143–53.

Cobo-Vuilleumier, N., and Gauthier, B.R. (2020). Time for a paradigm shift in treating type 1 diabetes mellitus: coupling inflammation to islet regeneration. Metabolism. 104: 154137.

Cobo-Vuilleumier, N., Lorenzo, P.I., Rodríguez, N.G., Herrera Gómez, I. de G., Fuente-Martin, E., López-Noriega, L., et al. (2018). LRH-1 agonism favours an immune-islet dialogue which protects against diabetes mellitus. Nat. Commun. 9: 1488.

Couri, C.E.B., Malmegrim, K.C.R., and Oliveira, M.C. (2018). New Horizons in the Treatment of Type 1 Diabetes: More Intense Immunosuppression and Beta Cell Replacement. Front. Immunol. 9: 1086.

Doyle, M.E., and Egan, J.M. (2007). Mechanisms of action of glucagon-like peptide 1 in the pancreas. Pharmacol. Ther. 113: 546–93.

Eleazu, C.O., Eleazu, K.C., Chukwuma, S., and Essien, U.N. (2013). Review of the mechanism of cell death resulting from streptozotocin challenge in experimental animals, its practical use and potential risk to humans. J. Diabetes Metab. Disord. 12: 60.

Engeli, S. (2008). Dysregulation of the endocannabinoid system in obesity. J. Neuroendocrinol. 20 Suppl 1: 110–5.

Forouhi, N.G., and Wareham, N.J. (2014). Epidemiology of diabetes. Medicine (Baltimore). 42: 698–702.

González-Mariscal, I., Krzysik-Walker, S.M., Doyle, M.E., Liu, Q.-R., Cimbro, R., Santa-Cruz Calvo, S., et al. (2016a). Human CB1 Receptor Isoforms, present in Hepatocytes and β-cells, are Involved in Regulating Metabolism. Sci. Rep. 6: 33302.

González-Mariscal, I., Krzysik-Walker, S.M., Kim, W., Rouse, M., and Egan, J.M. (2016b). Blockade of cannabinoid 1 receptor improves GLP-1R mediated insulin secretion in mice. Mol. Cell. Endocrinol. 423: 1–10.

González-Mariscal, I., Montoro, R.A., Doyle, M.E., Liu, Q.-R., Rouse, M., O’Connell, J.F., et al. (2018). Absence of cannabinoid 1 receptor in beta cells protects against high-fat/high-sugar diet-induced beta cell dysfunction and inflammation in murine islets. Diabetologia.

Hundhausen, C., Roth, A., Whalen, E., Chen, J., Schneider, A., Long, S.A., et al. (2016). Enhanced T cell responses to IL-6 in type 1 diabetes are associated with early clinical disease and increased IL-6 receptor expression. Sci. Transl. Med. 8: 356ra119.

International Diabetes Federation IDF Diabetes Atlas. Jourdan, T., Godlewski, G., Cinar, R., Bertola, A., Szanda, G., Liu, J., et al. (2013). Activation of the Nlrp3 inflammasome in infiltrating macrophages by endocannabinoids mediates beta cell loss in type 2 diabetes. Nat. Med. 19: 1132–40.

Katchan, V., David, P., and Shoenfeld, Y. (2016). Cannabinoids and autoimmune diseases: A systematic review. Autoimmun. Rev. 15: 513–528.

Kelleher, Z.T., Sha, Y., Foster, M.W., Foster, W.M., Forrester, M.T., and Marshall, H.E. (2014). Thioredoxin-mediated denitrosylation regulates cytokine-induced nuclear factor κB (NF-κB) activation. J. Biol. Chem. 289: 3066–72.

Lindley, S., Dayan, C.M., Bishop, A., Roep, B.O., Peakman, M., and Tree, T.I.M. (2005). Defective Suppressor Function in CD4+CD25+ T-Cells From Patients With Type 1 Diabetes. Diabetes 54: 92–99.

Maedler, K., Sergeev, P., Ris, F., Oberholzer, J., Joller-Jemelka, H.I., Spinas, G.A., et al. (2002). Glucose-induced beta cell production of IL-1beta contributes to glucotoxicity in human pancreatic islets. J. Clin. Invest. 110: 851–60.

McKillop, A.M., Moran, B.M., Abdel-Wahab, Y.H.A., and Flatt, P.R. (2013). Evaluation of the insulin releasing and antihyperglycaemic activities of GPR55 lipid agonists using clonal beta-cells, isolated pancreatic islets and mice. Br. J. Pharmacol. 170: 978–90.

McKillop, A.M., Moran, B.M., Abdel-Wahab, Y.H.A., Gormley, N.M., and Flatt, P.R. (2016). Metabolic effects of orally administered small-molecule agonists of GPR55 and GPR119 in multiple low-dose streptozotocin-induced diabetic and incretin-receptor-knockout mice. Diabetologia 59: 2674–2685.

Nir, T., Melton, D.A., and Dor, Y. (2007). Recovery from diabetes in mice by beta cell regeneration. J. Clin. Invest. 117: 2553–61.

Pacher, P., Kogan, N.M., and Mechoulam, R. (2020). Beyond THC and Endocannabinoids. Annu. Rev. Pharmacol. Toxicol. 60: annurev-pharmtox-010818-021441.

Romero-Zerbo, S.Y., García-Fernández, M., Espinosa-Jiménez, V., Pozo-Morales, M., Escamilla-Sánchez, A., Sánchez-Salido, L., et al. (2020). The Atypical Cannabinoid Abn-CBD Reduces Inflammation and Protects Liver, Pancreas, and Adipose Tissue in a Mouse Model of Prediabetes and Non-alcoholic Fatty Liver Disease. Front. Endocrinol. (Lausanne). 11:.

Ruz-Maldonado, I., Pingitore, A., Liu, B., Atanes, P., Huang, G.C., Baker, D., et al. (2018). LH-21 and abnormal cannabidiol improve β-cell function in isolated human and mouse islets through GPR55-dependent and -independent signalling. Diabetes, Obes. Metab. 20: 930–942.

Talaat, I.M., Nasr, A., Alsulaimani, A.A., Alghamdi, H., Alswat, K.A., Almalki, D.M., et al. (2016). Association between type 1, type 2 cytokines, diabetic autoantibodies and 25-hydroxyvitamin D in children with type 1 diabetes. J. Endocrinol. Invest. 39: 1425–1434.

Tomita, T. (2010). Immunocytochemical localization of cleaved caspase-3 in pancreatic islets from type 1 diabetic subjects. Islets 2: 24–9.

Tomko, A., O’Leary, L., Trask, H., Achenbach, J.C., Hall, S.R., Goralski, K.B., et al. (2019). Antitumor Activity of Abnormal Cannabidiol and Its Analog O-1602 in Taxol-Resistant Preclinical Models of Breast Cancer. Front. Pharmacol. 10: 1124.

Vong, C.T., Tseng, H.H.L., Kwan, Y.W., Lee, S.M.-Y., and Hoi, M.P.M. (2019a). G-protein coupled receptor 55 agonists increase insulin secretion through inositol trisphosphate-mediated calcium release in pancreatic β-cells. Eur. J. Pharmacol. 854: 372–379.

Vong, C.T., Tseng, H.H.L., Kwan, Y.W., Lee, S.M.-Y., and Hoi, M.P.M. (2019b). Novel protective effect of O-1602 and abnormal cannabidiol, GPR55 agonists, on ER stress-induced apoptosis in pancreatic β-cells. Biomed. Pharmacother. 111: 1176–1186.

Weiss, L., Zeira, M., Reich, S., Slavin, S., Raz, I., Mechoulam, R., et al. (2008). Cannabidiol arrests onset of autoimmune diabetes in NOD mice. Neuropharmacology 54: 244–249.

Yao, Y., Xu, X.H., and Jin, L. (2019). Macrophage polarization in physiological and pathological pregnancy. Front. Immunol. 10: 792.

Zhou, R., Tardivel, A., Thorens, B., Choi, I., and Tschopp, J. (2010). Thioredoxin-interacting protein links oxidative stress to inflammasome activation. Nat. Immunol. 11: 136–140.

